## Supplemental Figs1-3 and Table S1 for "Plasticity of gene expression in the nervous system by exposure to environmental odorants that inhibit HDACs"

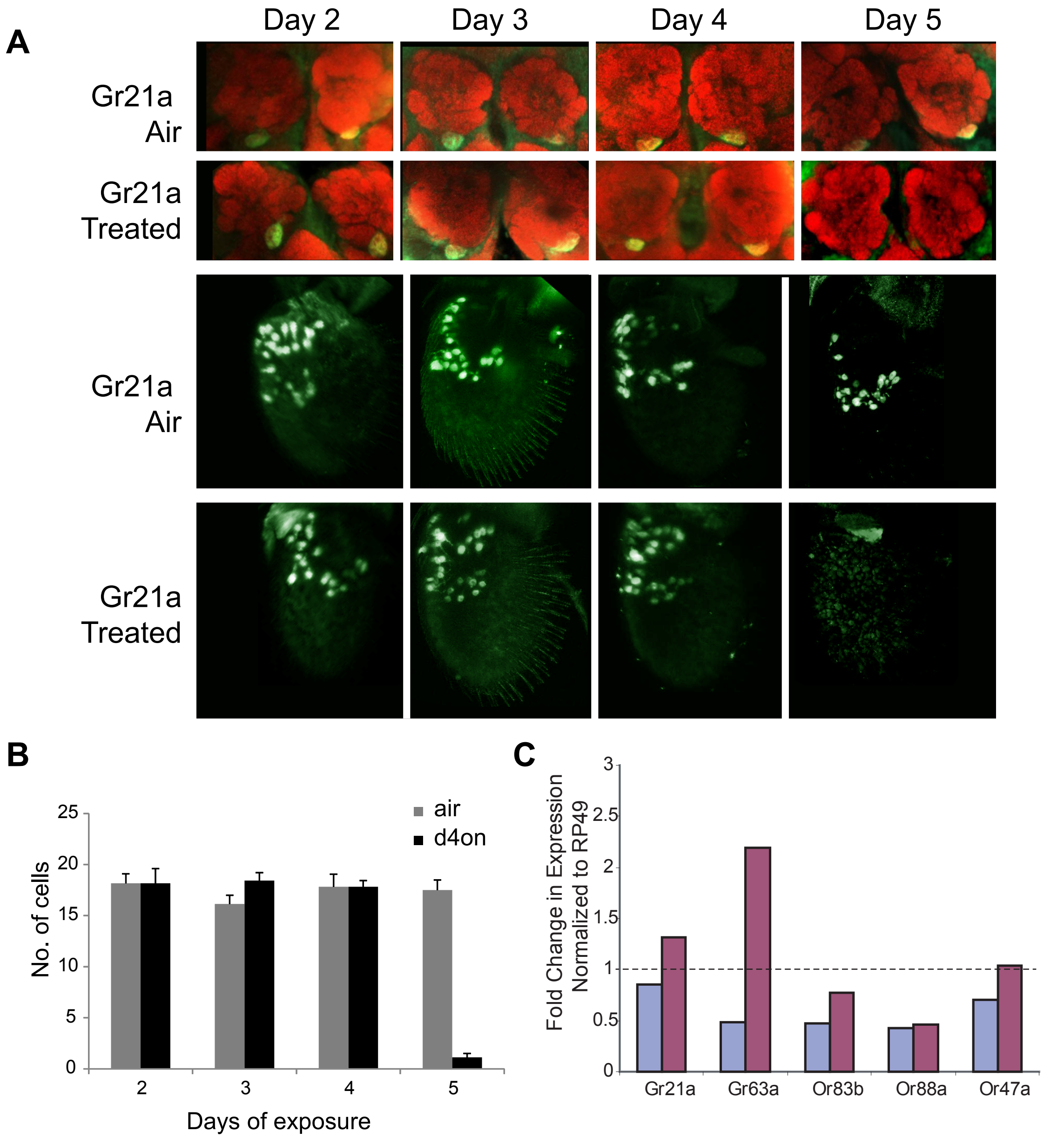


**Fig. S1. $CO_2$ inhibitory odor diacetyl causes down regulation of $CO_2$ receptor. (A)** Antennal and whole mount brain staining of *Gr21a-Gal4* flies. Flies were exposed to headspace from 1% diacetyl (v/v in paraffin oil) or air for 2 to 5 days (see methods). Brains and antenna were dissected on the indicated days, fixed, and then stained for neuropil marker nc82 (red) and anti-GFP (green). **(B)** Mean of ab1C neurons expressing GFP after indicated days of odor exposure. d4on=diacetyl. n=6, error bars=s.e.m. **(C)** Flies were exposed to headspace from 1% diacetyl for 6 days (Day 6 treated, blue bars), or exposed to headspace from 1% diacetyl for 6 days and allowed to recovery in air for 5 days (Day 6 treated, 5-day recovery, purple bars). Gene expression by QPCR is compared to flies raised in air for the equivalent numberof days. Expression data is based on fold expression normalized to RP49. Fold expression=1 indicates no change in expression. n=2.

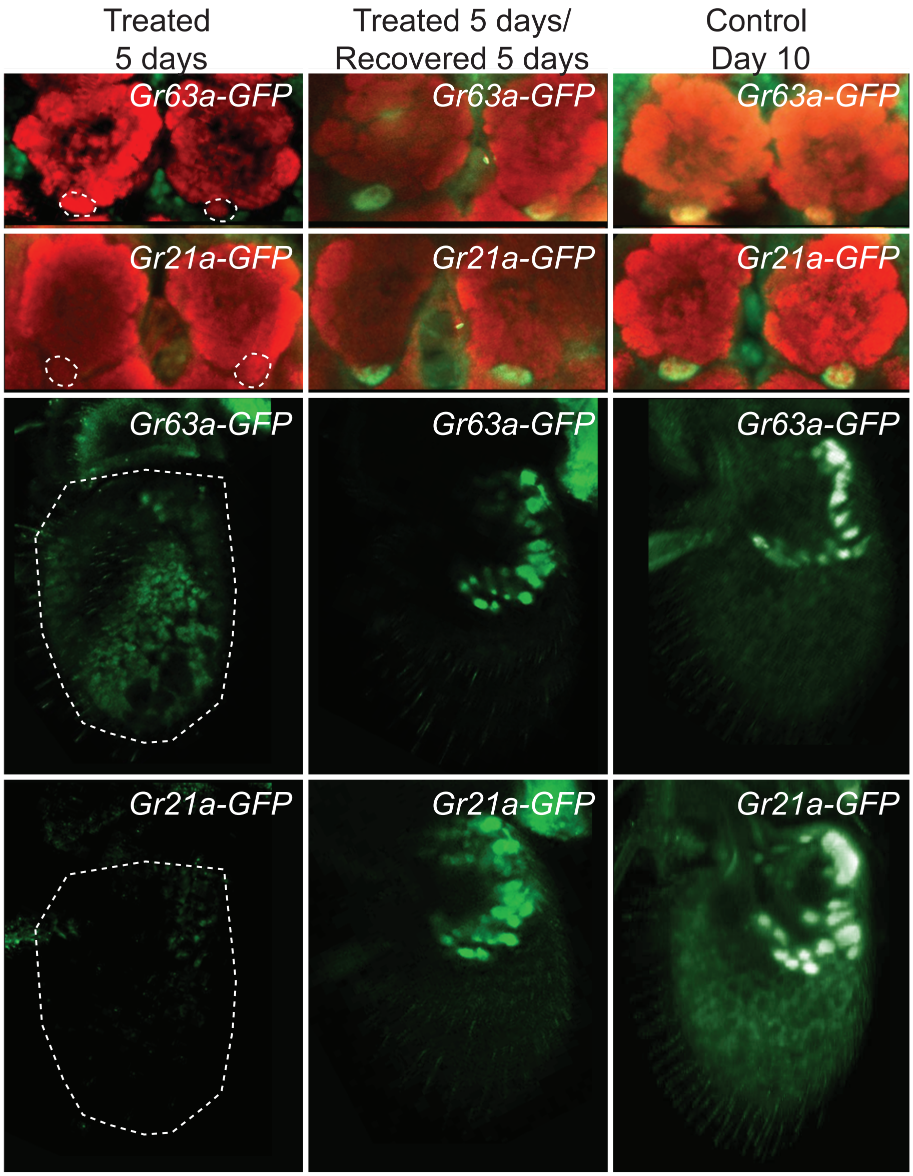


**Fig. S2. Down regulation of the $CO_2$ receptor driver is reversible.**

Representative images of antennal and whole mount brain staining of *Gr63a-Gal4*;*UAS-mcd8GFP* and *Gr21a-Gal4; UAS-mcd8GFP* flies. Flies were exposed to headspace from 1% diacetyl (v/v in paraffin oil) for 5 days (treated 5 days) and allowed to recover in clean air for 5 days (treated 5 days/recovered 5 days) or exposed to air for 10 days (control day 10). Brains and antenna were dissected on the indicated days, fixed, and then stained for neuropil marker nc82 (red) and anti-GFP (green).

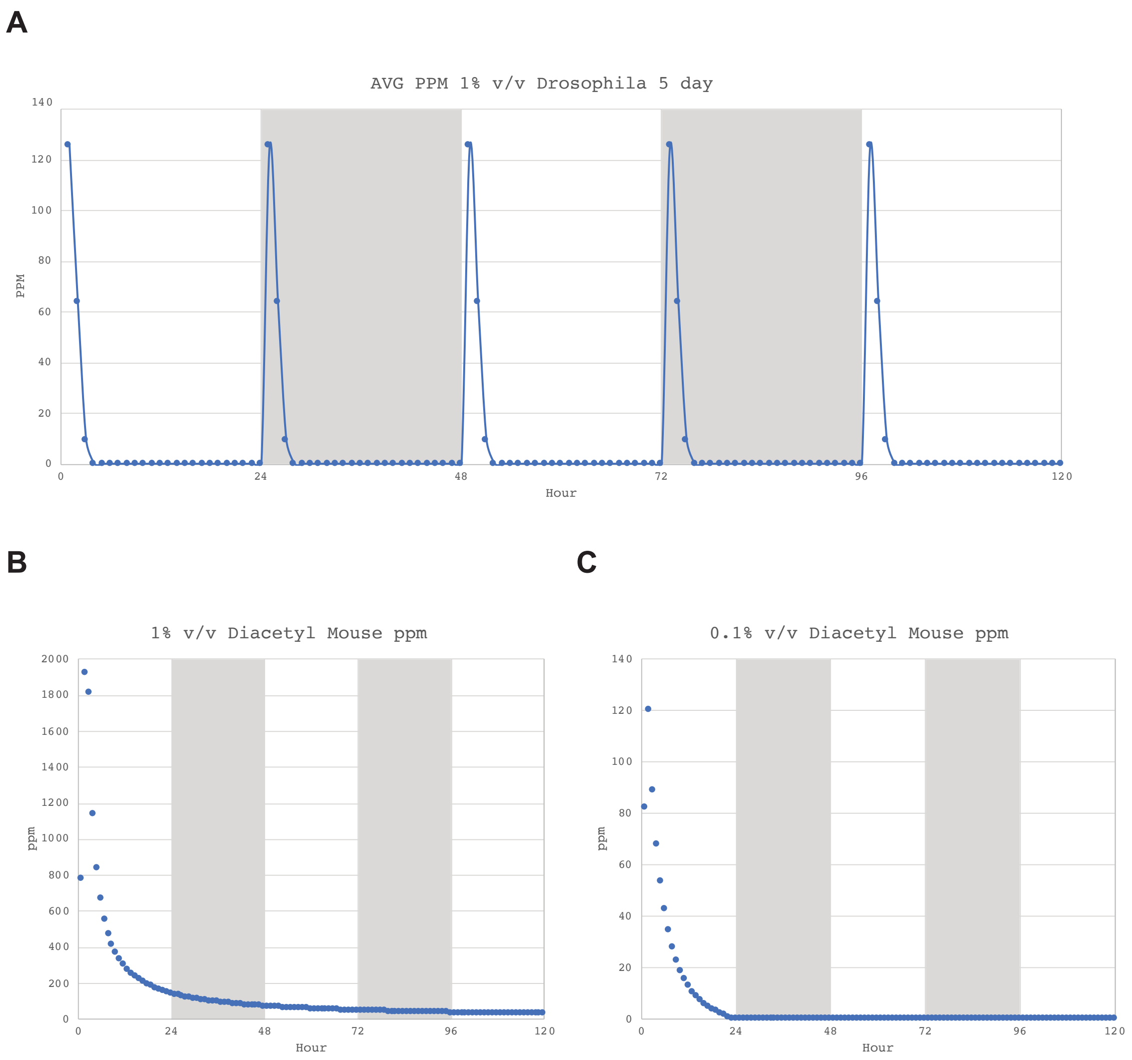


**Fig. S3. Dose of diacetyl in ppm in experimental chambers over time**

Mean concentration of diacetyl in the air in the exposure experimental chambers in ppm calculated based on weight loss of the compound.

**TABLE S1.** Concentration of diacetyl in some common sources.

| # | Common sources of Diacetyl | AVG Diacetyl (ppm) | References |
|---|---|---|---|
| 1 | Starter Distillates (SDL) in Dairy Product Production | 1.2-22,000 | Rincon-Delgadillo, M. I., Lopez-Hernandez, A., Wijaya, I., & Rankin, S. A. (2012). Diacetyl levels and volatile profiles of commercial starter distillates and selected dairy foods. *Journal of dairy science*, *95*(3), 1128-1139. |
| 2 | Mainstream cigarette smoke | 250-361 | Pierce, J.S., et al., *Diacetyl and 2,3-pentanedione exposures associated with cigarette smoking: implications for risk assessment of food and flavoring workers.* Crit Rev Toxicol, 2014. **44**(5): p. 420-35. |
| 3 | Microwave Popcorn Facility | 1-57.2 | Kanwal, R., et al., *Occupational Lung Disease Risk and Exposure to Butter-Flavoring Chemicals After Implementation of Controls at a Microwave Popcorn Plant.* Public Health Reports, 2011. **126**(4): p. 480-494. |
| 4 | Baked goods | 44 | Hall, R.L. and B.L. Oser, *Recent Progress in Consideration of Flavoring Ingredients under Food Additives Amendment .3. Gras Substances.* Food Technology, 1965. **19**(2p2): p. 151-&. |
| 5 | Candy | 21-35 | Hall, R.L. and B.L. Oser, *Recent Progress in Consideration of Flavoring Ingredients under Food Additives Amendment .3. Gras Substances.* Food Technology, 1965. **19**(2p2): p. 151-&. |
| 6 | Gelatins/Puddings | 19 | Hall, R.L. and B.L. Oser, *Recent Progress in Consideration of Flavoring Ingredients under Food Additives Amendment .3. Gras Substances.* Food Technology, 1965. **19**(2p2): p. 151-&. |
| 7 | Unflavored Brewed cup of Coffee | 7 | Pierce, J.S., et al., *Characterization of naturally occurring airborne diacetyl concentrations associated with the preparation and consumption of unflavored coffee.* Toxicol Rep, 2015. **2**: p. 1200-1208.<br><br>Yeretzian, C., A. Jordan, and W. Lindinger, *Analysing the headspace of coffee by proton-transfer-reaction mass-spectrometry.* International Journal of Mass Spectrometry, 2003. **223**(1-3): p. 115-139. |
